# Homogeneous inhibition is optimal for the phase precession of place cells in the CA1 field

**DOI:** 10.1101/2022.04.24.489307

**Authors:** G.K. Vandyshev, I.E. Mysin

**Author notes:** Email addresses* (Vandyshev G.K.), (Mysin I.E.).

## Abstract

Place cells are hippocampal neurons encoding the position of an animal in space. Studies of place cells are essential to understanding the processing and transmission of information by neural networks of the brain. An important characteristic of place cell spike trains is phase precession. When an animal is running through the place field, the discharges of the place cells shift from the ascending phase of the theta rhythm through the minimum to the descending phase. The role of excitatory inputs to CA1 pyramidal neurons along the Schaffer collaterals and the perforant pathway in phase precession is described, but the role of inputs from local interneurons to CA1 pyramidal cell is poorly understood. We have set the goal the contribution of field CA1 interneurons to the phase precession of place cells using mathematical methods. The CA1 field was chosen because it provides the largest set of experimental data required to build and verify the model. We have solved the optimization problem and found the parameters of the excitatory and inhibitory inputs to the pyramidal neuron of the CA1 field so that the neuron generates a spike train with the effect of phase precession. We have discovered that the uniform inhibition of field CA1 pyramidal neurons best explains the effect of phase precession. Among interneurons, axo-axonal neurons make the greatest contribution to the inhibition of pyramidal cells.

## 1. Introduction

Place cells are the principal hippocampal neurons encoding space in mammals [1, 2, 3]. Although hippocampal neurons can encode time, smells, novelty, or images of people [4, 5, 6], spatial memory is the most investigated aspect. Place cells are convenient to examine for several reasons. First, it is easy to simultaneously observe the activity of neurons and the position of an animal in space. Secondly, about 50% of the principal neurons of the hippocampus exhibit the property of place cells in one maze [7, 8], so the use of even a few electrodes makes it possible to register place cells. The combination of fundamental importance and convenience in investigation has made place cells a model object in neuroscience, like Drosophila fly in genetics or E. coli in microbiology.

Place cells encoding places close to each other are combined into neural ensembles [8, 9]. Information processing and transmission between regions of the hippocampus is coordinated with gamma (25-120 Hz) and theta (4-12 Hz) rhythms [2, 10, 11]. Almost all neurons in the hippocampus and entorhinal cortex (EC) are in a theta rhythm phase when they are most likely to fire [10, 12, 13]. For example, parvalbumin (PV) basket interneurons fire in the descending phase, while OLM interneurons fire at the trough of the theta cycle [12, 14, 15]. Here and further, we will refer to the theta rhythm phase in the pyramidal layer of the CA1 field.

Pyramidal neurons of the CA1 field outside their place field discharge with rare spikes mainly at the minimum of the theta cycle [10, 12]; however, when an animal is in the place field, their spike rate increases, and the nature of their activity coupling to the theta phase the rhythm changes. When an animal runs through a place field, the place cells start to fire in the ascending phase of the theta rhythm. Further, their spikes shift through the minimum to the descending phase. Interspike interval is slightly less than the theta cycle. This phenomenon is called phase precession [1, 16]. The phenomenon of phase precession shows the complex structure of inputs to the pyramidal neuron while an animal is running through the place field.

Raising the spike rate in the place field indicates increased excitation and/or reduced inhibition of the pyramidal neuron. Studies of excitatory inputs into field CA1 show that excitation of the place cell when an animal is running into the place field is provided by the input from the EC, while when running out, the main role in maintaining firing activity passes to the input from the CA3 field [17, 18]. With such an excitation structure, phase precession is formed due to the moiré effect from the superposition of two phase-shifted oscillator inputs, resulting in oscillations of a higher frequency than the incoming signals have. Although there are other hypotheses to explain the displacement of place cell discharges in theta phase, the theory of two inputs has received the most experimental support [1, 17, 18, 19, 20, 21].

The role of inhibition in the formation of place field and phase precession, on the contrary, is unclear. On the one hand, optogenetic inhibition of PV neurons weakens the effect of phase precession [14]. PV interneurons fire predominantly when an animal is running into the place field, while the firing rate of OLM neurons, on the contrary, increases when it is running out of the place field [17]. Moreover, a recent study shows that not only PV and OLM, but also other types of interneurons have spatial modulation [22]. All these data support the idea that interneurons inhibit pyramidal cells weaker in the place field, and stronger outside the place field. Due to this reason, interneurons contribute to the selection of an active neural ensemble and the formation of phase precession.

On the other hand, there is evidence that contradicts this hypothesis. Grienberger recorded intracellular potentials of pyramidal cells and interneurons in freely-behaving animals [20]. The authors found no differences in the inhibitory effect on pyramidal neurons in the place field and outside the place field. According to these data, the discharges of pyramidal cells in place field occur only due to the amplification of excitatory inputs. Nevertheless, the authors emphasize that their data do not refute the idea of the participation of interneurons in the formation of phase precession. The contribution of interneurons to the formation of phase precession is possible due to the inhibition of pyramids in different phases of the theta rhythm [20].

We have applied a modeling approach to find an answer to the following question: what structure of excitation and inhibition of pyramidal neurons is optimal for the occurrence of phase precession based on the given phase relationships between external inputs (CA3 field and EC) and local interneurons of the CA1 field? From a mathematical point of view, we have solved the optimization problem. We have found the optimal input parameters for the best match between the model and the experimental data on phase precession.

## 2. Methods

### Pyramid neuron model

We used the pyramidal neuron model proposed by Ferguson and Campbell [23]. The neuron model has two compartments: perisomatic zone and apical dendrite. The equations for the soma and dendrite potentials are

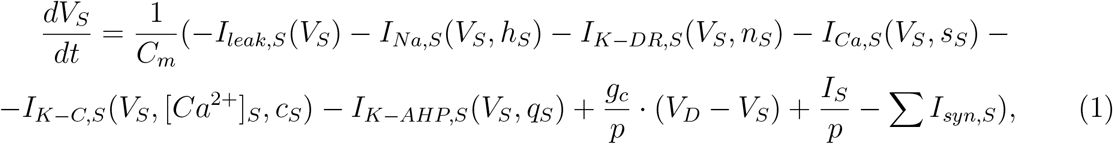

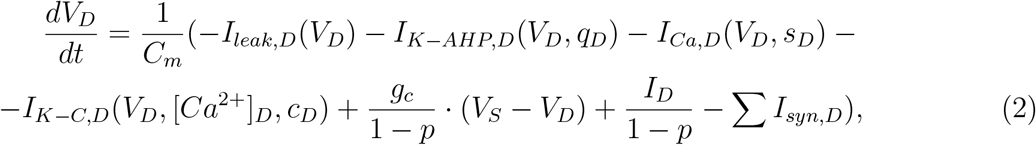

*j* ∈ {*S, D*}, *S* and *D* denote the soma and dendrite of pyramidal neurons, respectively. The equations for the currents and description of the parameters are given in Appendix

### The inputs model

The synaptic currents were given by the equation:

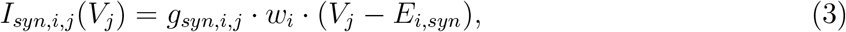

where *j* ∈ {*S, D*}, *S* and *D* denote input on the soma and dendrite; *E_syn_* is the reversal potential of the synaptic currents. *i* denote input type, *i* ∈{ca3, ec3, cck, pv, ngf, ivy, olm, aac, bis}; inputs data see in Table 1. Synaptic conductivities were described by double exponential equation:

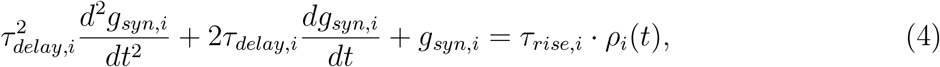

where *ρ_i_*(*t*) is a spike rate of presynaptic populations.

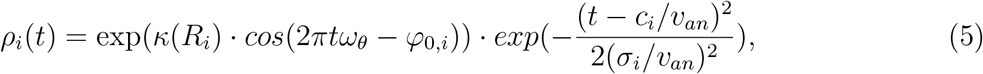

**Table 1:**
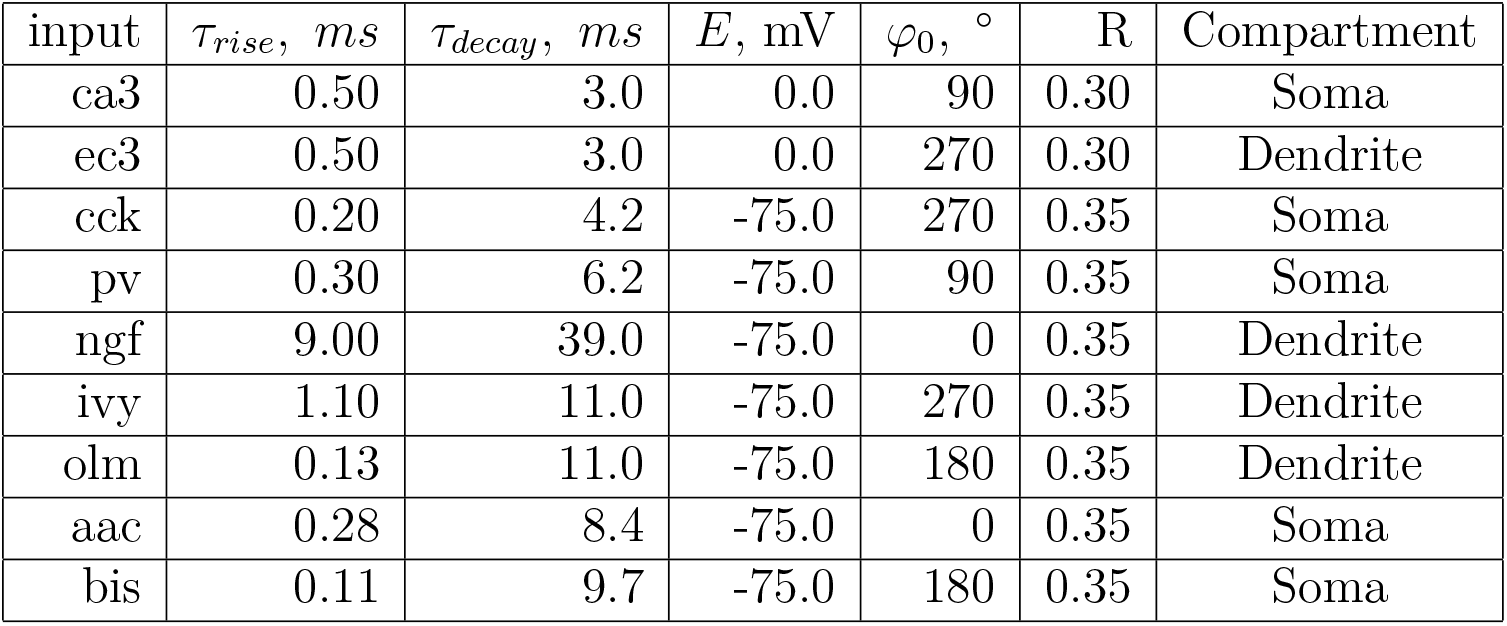
Input’s parameters

*v_an_* is animal velocity (cm/sec), *τ_rise,i_*, *τ_decay,i_, E_syn,i_, φ*_0,*i*_, *R_i_* are input’s parameters (Table 1); *w_i_*, *c_i_* (cm), *σ_i_* (cm) are parameters for optimization. *R_i_* is ray length, degree of neuron firing coupling to a phase of rhythm. *κ* can be calculated from *R* by following equations:

Approximation of *κ* from *R* [24]

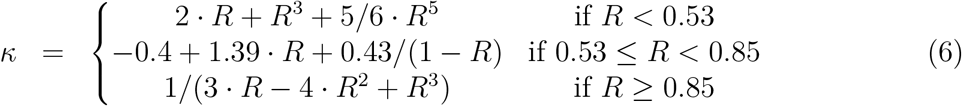

### Optimization of the model

We used a differential evolution algorithm to minimize the loss function [25, 26]. We optimize the model parameters so that the spike rate of the pyramidal neuron coincides with the target one. Simulated firing rate is a smoothed spike train:

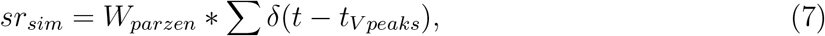

where *sr_sim_* is a simulated spike rate; * is convolution, *W_parzen_* is a Parzen window (number points = 1001); *t_V_peaks__* is times of spikes. Spikes are local maxima of somatic potential greater than −10 mV. Theta phases are:

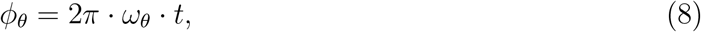

The target spike rate was given by the following equation:

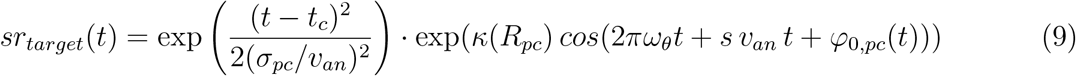

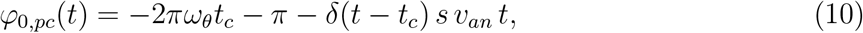

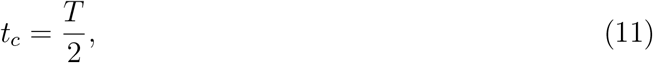

where *s* is a precession slope (*rad/cm*); *ω*_Θ_ is the frequency of theta rhythm (Hz); *t_c_* is the center of the place field, in our simulation *t_c_* is half of the period of simulation (T). *σ_pc_* is the size of place field; *R_pc_* is the strength of phase coupling of place cell to theta phase. Loss function:

Loss function equation

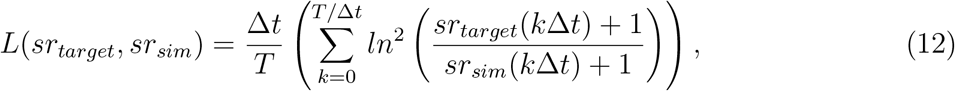

Δ*t* = 0.1 ms is an integration step. Parameters for optimization algorithm: population size = 24; absolute tolerance = 0.001; recombination = 0.7; mutation = 0.2; strategy is “best2bin”.

Parameters for basic model: precession slope *s* = 5°/cm; animal velocity *ν_an_* = 20 cm/sec; *R_pc_* = 0.5, *σ_pc_* = 4 cm, theta frequency *ω_θ_* = 8 Hz. Please note: in the equations, the phases are expressed in radians, but we used degrees on the plots and for source data for better visual representation.

### Statistical processing of the results

The analysis of phase precession was carried out as is customary in experimental works. The slope of the phase precession was estimated using circular-linear regression [27]. The strength of the relationship between the animal’s position and the theta rhythm phase was evaluated using a circular-linear correlation [28, 29].

### Software tools and implementation

We coded simulations in Python 3.9 with standard scientific packages Numpy, Scipy, and Matplotlib [30, 31, 32]. The Cython and Numba packages were used to speed up computations [33, 34]. To solve the optimization problem, the “*differential_evolution*” function from the Scipy package was applied. Our code is available on [GitHub](https://github.com/ivanmysin/Phase_precession).

## 3. Results

### Basic model

The pyramidal neuron of the CA1 field receives two excitatory inputs: from the CA3 field and 3-th layer of the EC, and many inhibitory inputs from local interneurons (Fig. 1). We consider inputs from 7 types of interneurons: PV and CCK basket cells, OLM, Ivy, neurogliaform (NGF), bistratified (BIS), and axo-axonal (AAC) cells. The interneurons of these types are the most studied and account for more than 70% of all interneurons in the CA1 field [35].

**Figure 1:**
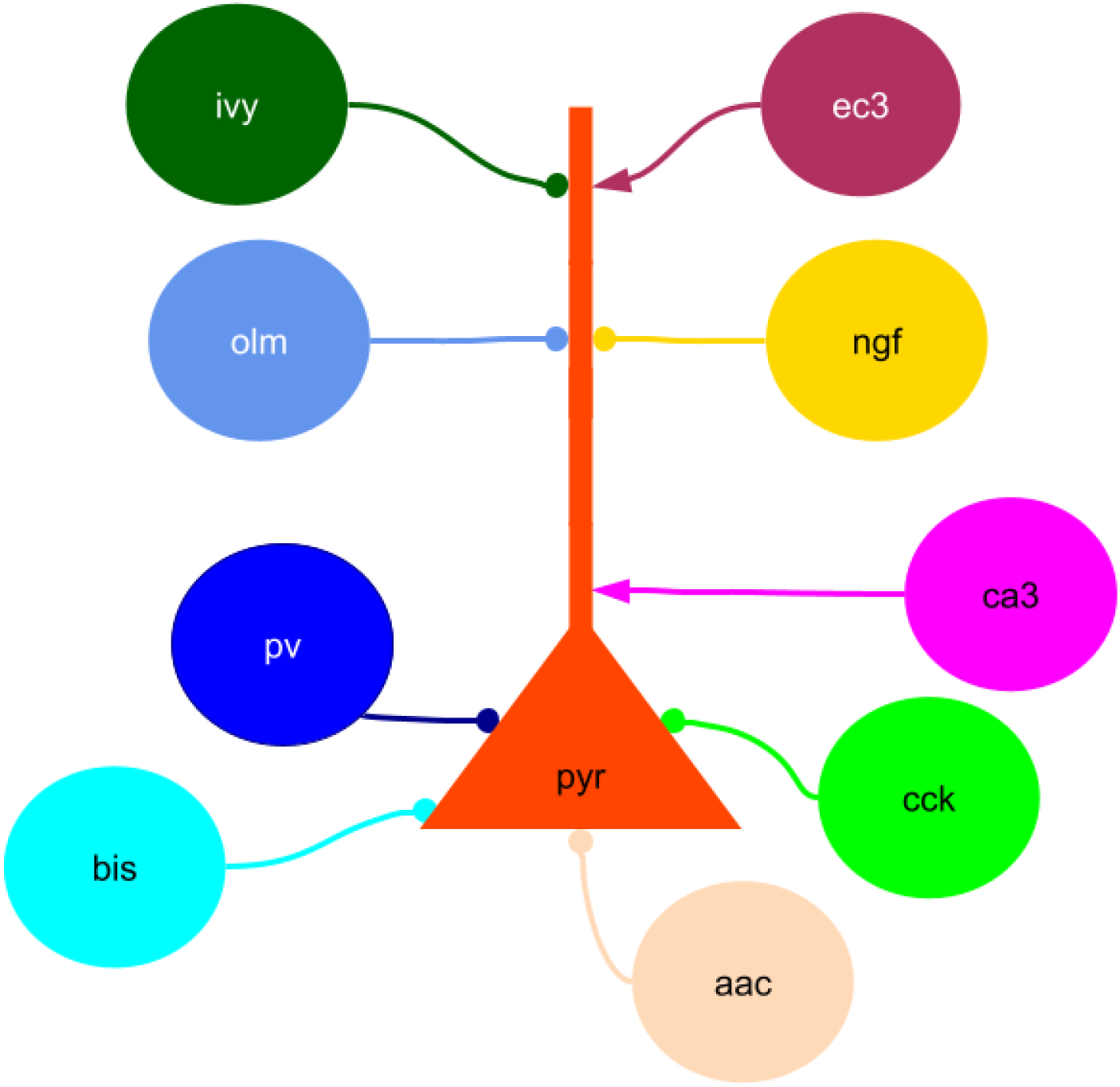
Scheme of the model. Neuron notations: pyr - pyramidal cells, pv - PV basket cells, olm - Oriens-lacunosum molecular cells, cck - CCK basket cells, ivy - Ivy cells, ngf - neurogliaform cells, bis - bistratified cells, aac - axo-axonal cells.

We consider the case when an animal runs along a linear track (Fig. 2). All inputs were modeled by functions (eq. 3–5). The input approximation assumes that each input is generated by a large population of presynaptic cells. Each presynaptic population has a coupling to the phase of theta rhythm and spatial modulation (Table 1). Spatial modulation of the inputs (*σ* from eq. 5) consists of spatial modulation of individual neurons and the distribution of connections. The spatial modulation of the input is small even in the case of strong spatial modulation of all neurons of the presynaptic population if all connections to the pyramidal neuron have equal synaptic weight. In other words, if presynaptic neurons encoding distant places give the same input to the one pyramidal cell, then the total synaptic conductivity is not spatially modulated.

**Figure 2:**
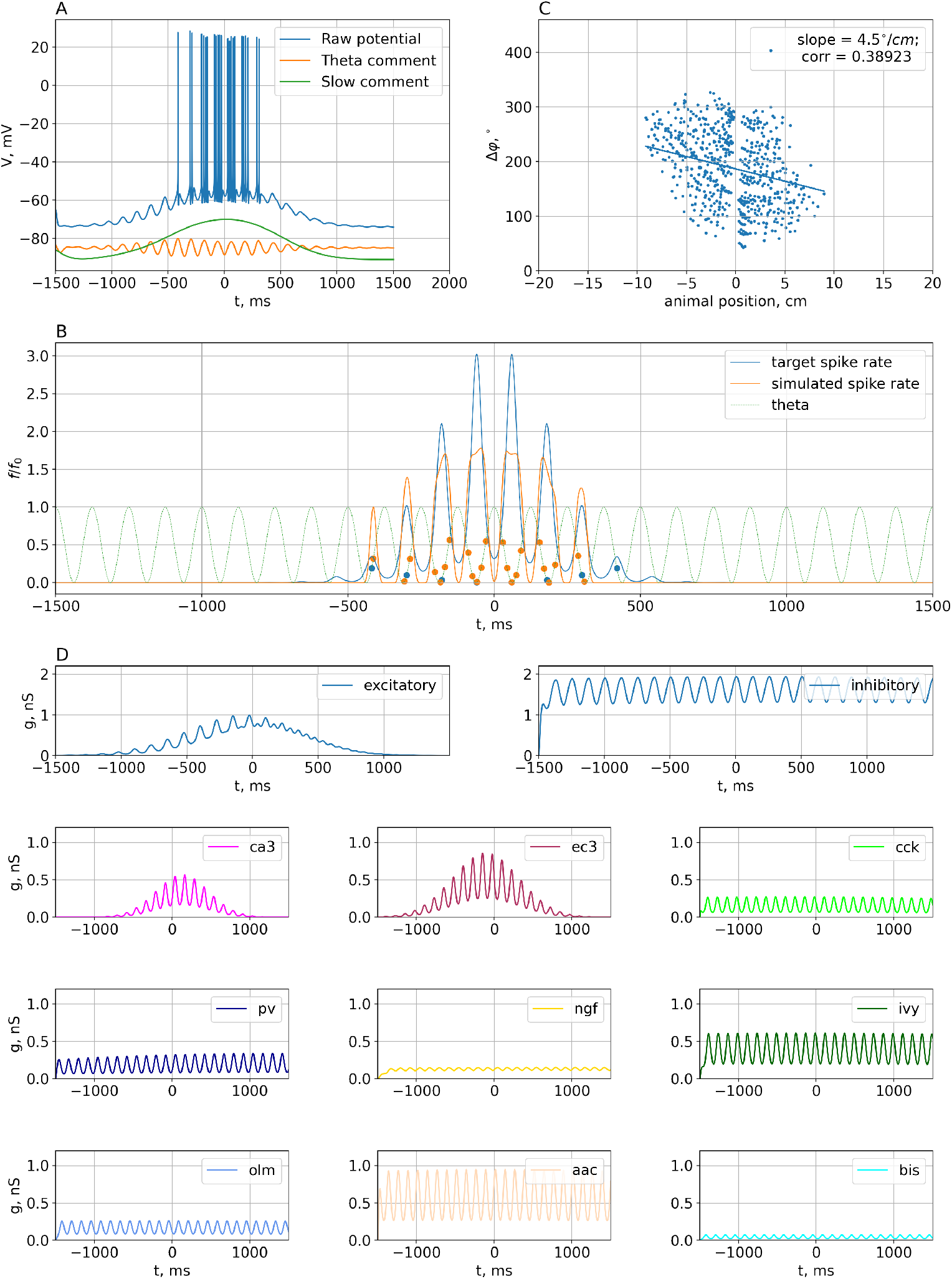
Results of optimization of the basic model. A. Dynamics of intracellular potential on the soma of the pyramidal neuron. B. Target spike rate (blue) (equation 7) and spike rate of the optimized model (orange) (equation 9). C. Dependence of firings of the pyramidal neuron from the position of the animal. The plot is based on a series of simulations. The simulations differed in the frequency of the theta rhythm (from 4 to 12 Hz with a step of 2 Hz) and the speed of the animal (from 10 to 30 cm/sec with a step of 5). D. Optimal conductivities. The upper plots show the sums of excitatory and inhibitory conductivities. The lower plots show individual conductivities for each input.

Non-optimized parameters of the loss function (for example, the slope of phase precession) or the input function (ray length, theta phase) were selected according to the mean and median values measured in the experiments [17, 36]. To verify the accuracy of the solution, we have simulated a model with the optimized parameters, but with different theta rhythm frequencies and the speed of movement of the animal (Fig. 2C).

We have found the optimal configuration of excitatory and inhibitory inputs to the CA1 pyramidal neuron to explain the effect of phase precession. Fig. 2 shows the results of optimization of the parameters of the inputs to reproduce the phase precession. Phase precession occurs as a result of excitatory inputs. When an animal is running into the place field, excitation from the input from the EC dominates, and when it is running out, the input from the CA3 field does (Fig. 2D). This is consistent with experimental data [17] and well described by computational models [19, 37].

The optimization results show that homogeneous inhibition best explains the effect of phase precession (Fig. 2D). Parameter *σ* (eq. 5) of inputs from all types of interneurons is much larger than the place field of the pyramidal neuron. With these place field values, the position of the center relative to the center of the place field (the value of c in equation 5) is not significant.

To make sure the result is reliable, we have run 30 optimizations to random sets of non-optimized parameters. The algorithm convergences to similar results in all cases (Fig. 3). All groups of interneurons had large sigmas in most simulations. Some cases of small *σ* correspond to small weights or distant centers, which excludes their significant influence on the formation of phase precession. Note that *σ* of the exciting inputs (mainly EC3) in several simulations were also large. In these cases, the spatial modulation of the pyramid cell was provided by only one input, but the mechanism of phase precession as a result of the moiré effect is not changed.

**Figure 3:**
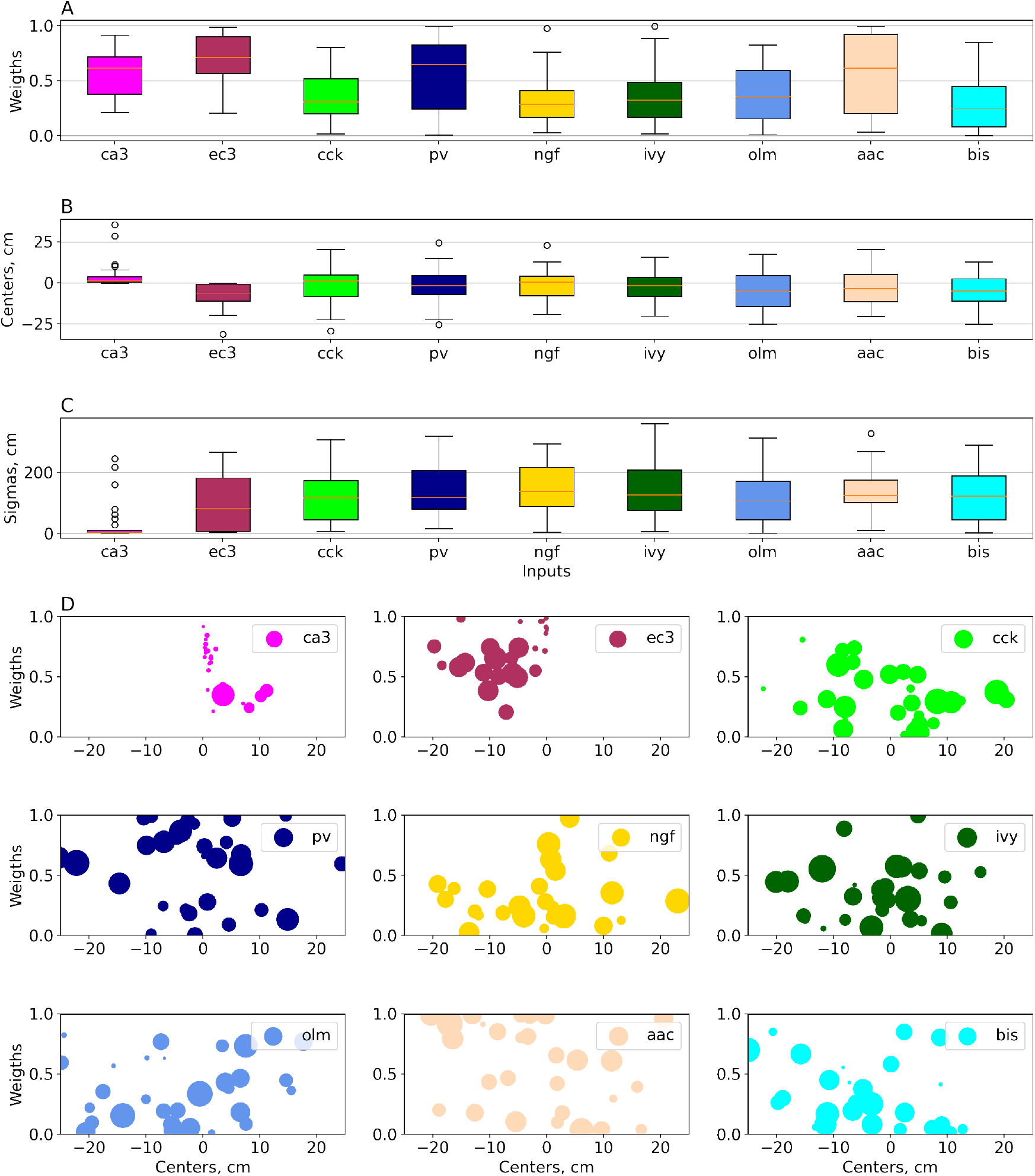
Optimization results for random sets of parameters. In each optimization, the value of following parameters were randomly selected from a uniform distribution: theta rhythm frequency (range 4-12 Hz), phase precession slope (2.5 - 7 deg/cm), animal movement speed (10 - 30 cm/sec), place field size (sigma, 2 - 5 cm). 30 sets of parameters have been studied. A. Distribution of weights. B. Distribution of centers. C. Distribution of the size of place fields (sigma). D. Scatter plot for same data as in A, B, C, size of dots is proportional to sigma. Large points correspond to homogeneous inputs.

### Sensitivity of the model to optimized parameters

We investigated the sensitivity of the phase precession to optimized parameters: centers, sizes, and weights of the inputs. Despite the uniformity of inputs from all interneurons, their weights differ significantly. Fig. 3A demonstrates that axo-axonal cells have the most weight. This is because axo-axonal cells are active at the peak of the theta rhythm; when firings of pyramidal neurons are minimal [12, 13]. The weight of bistratified cells, on the contrary, is minimal; since their firings occur at the minimum of the theta rhythm when pyramidal neurons are the most excited. Spikes of OLM neurons also occur at the minimum of the theta rhythm, but they inhibit the apical dendrite. When running out of the place field, the dominant input falls on the perisomatic zone of pyramidal cells; therefore, inhibition from OLM neurons has an insignificant effect on phase precession. The weights of the remaining interneurons have an intermediate value, which is necessary to prevent “false” firings of pyramidal neurons. When the weights of PV and NGF neurons performing inhibition in the descending phase of the theta rhythm were set to zero, the pyramidal neuron slightly increased activity at the initial stage of the trajectory. Correspondingly, when inputs inhibiting the ascending phase of the theta rhythm (CCK, Ivy) were excluded from the model, the pyramidal neuron increased its activity upon the input from the place field (Fig. 4). Thus, inhibition, although it does not have spatial modulation, it makes a significant contribution to the formation of phase precession due to the distribution of the theta rhythm phases.

**Figure 4:**
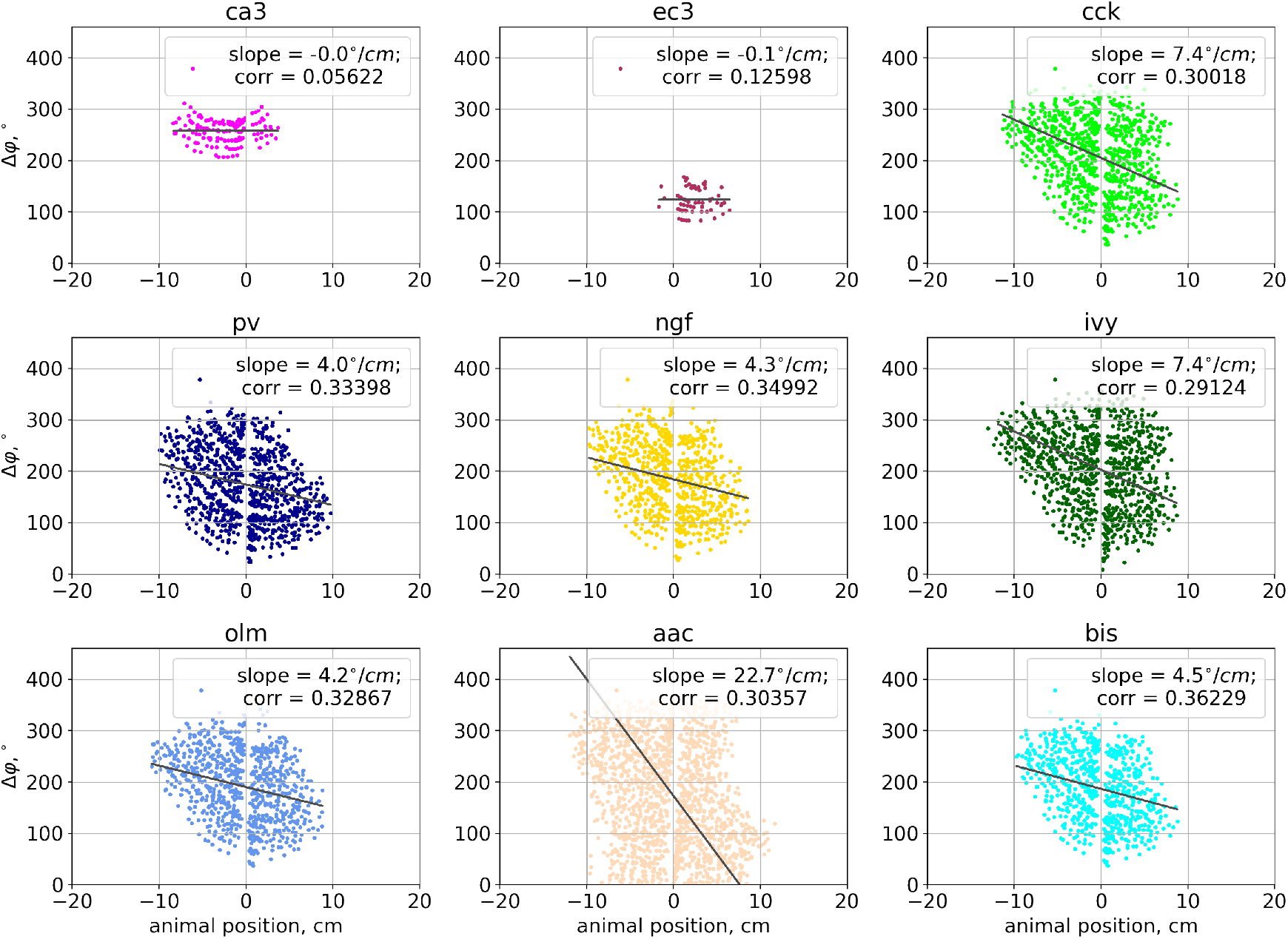
Estimation of the significance of the weight of each input. Phase precession at zero weight of each input.

To assess the possibility of the existence of a configuration with non-uniform inhibition, we have run an optimization with a constraint on *σ* in equation (5) (Fig. 5A, B, C). Interneurons except for axo-axonal cells have small weights or distant centers, which excludes their significant influence on the formation of phase precession. To ensure the reliability of this result, we have performed 30 optimizations in different sets of non-optimized parameters. The algorithm converges to similar values in most simulations, but in several cases we found opposite results. In 10 simulations, the input from bistratified neurons had a large weight (*w* > 0.5) and a small distance to the center of the place field (−4 cm < *c* <0 cm). In this case, the activity of bistratified cells do not allow spikes of pyramidal neuron, which can be produced by the input from the CA3 field, when the animal is running into the place field. There are several cases of large weights and close centers, they probably correspond to the local minima of the optimization function, so we don’t discuss them.

**Figure 5:**
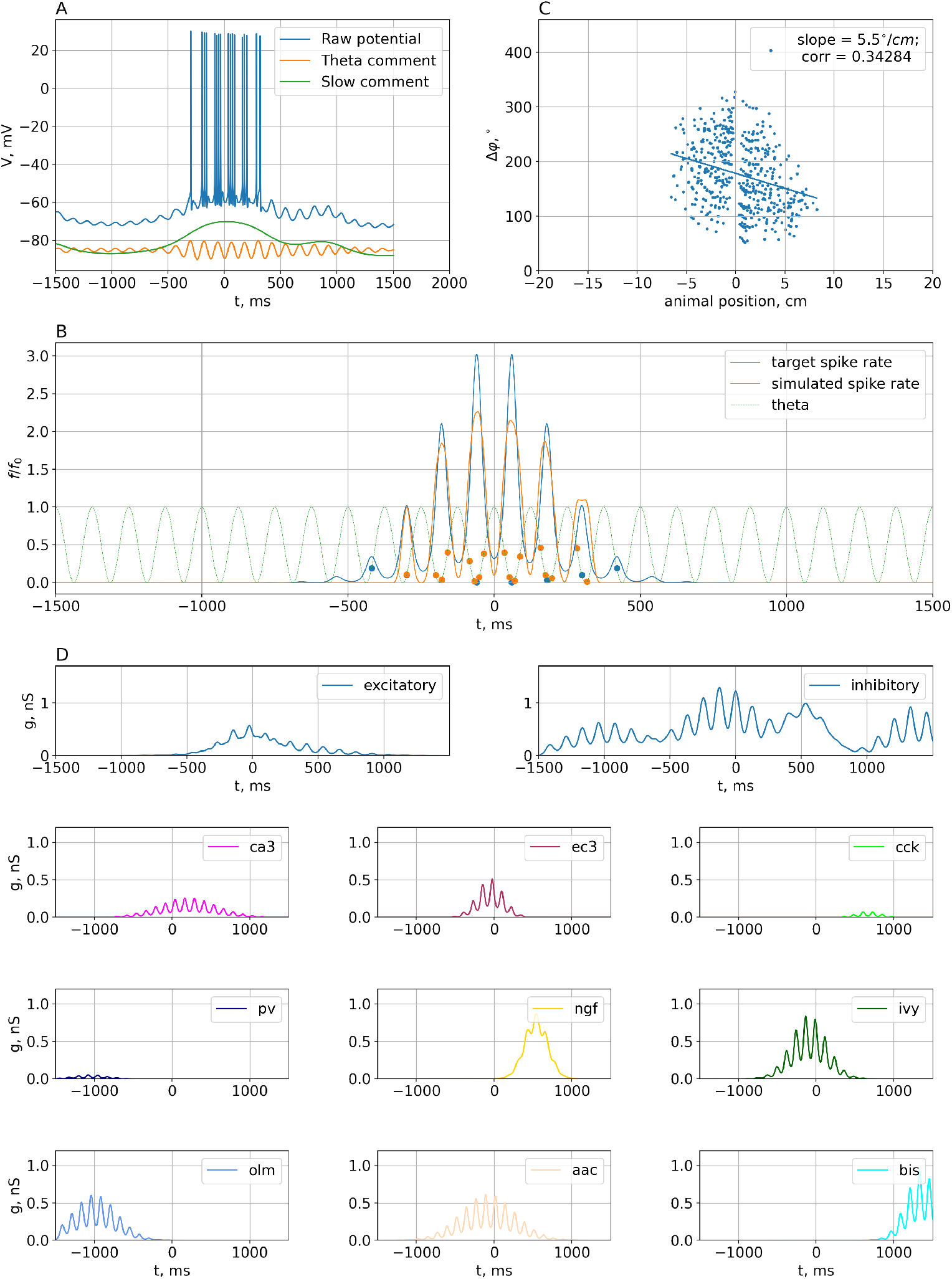
Estimation of the significance of spatial modulation of inputs. The spatial modulation of the inputs is limited, *σ* is no more than 8 cm. The same parameters as in the base model were taken. The drawing is similar to Fig. 2. A. Dynamics of intracellular potential on the soma of a pyramidal neuron. B. Target spike rate (blue) (equation 7) and spike rate of the optimized model (orange) (equation 9). C. Dependence of firings of a pyramidal neuron from the position of the animal. The plot is based on a series of simulations. The simulations differed in the frequency of the theta rhythm (from 4 to 12 Hz with a step of 2 Hz) and the speed of the animal (from 5 to 30 cm/sec with a step of 5). D. Optimal conductivities. The upper plots show the sums of excitatory and inhibitory conductivities. The lower plots show individual conductivities for each input.

**Figure 6:**
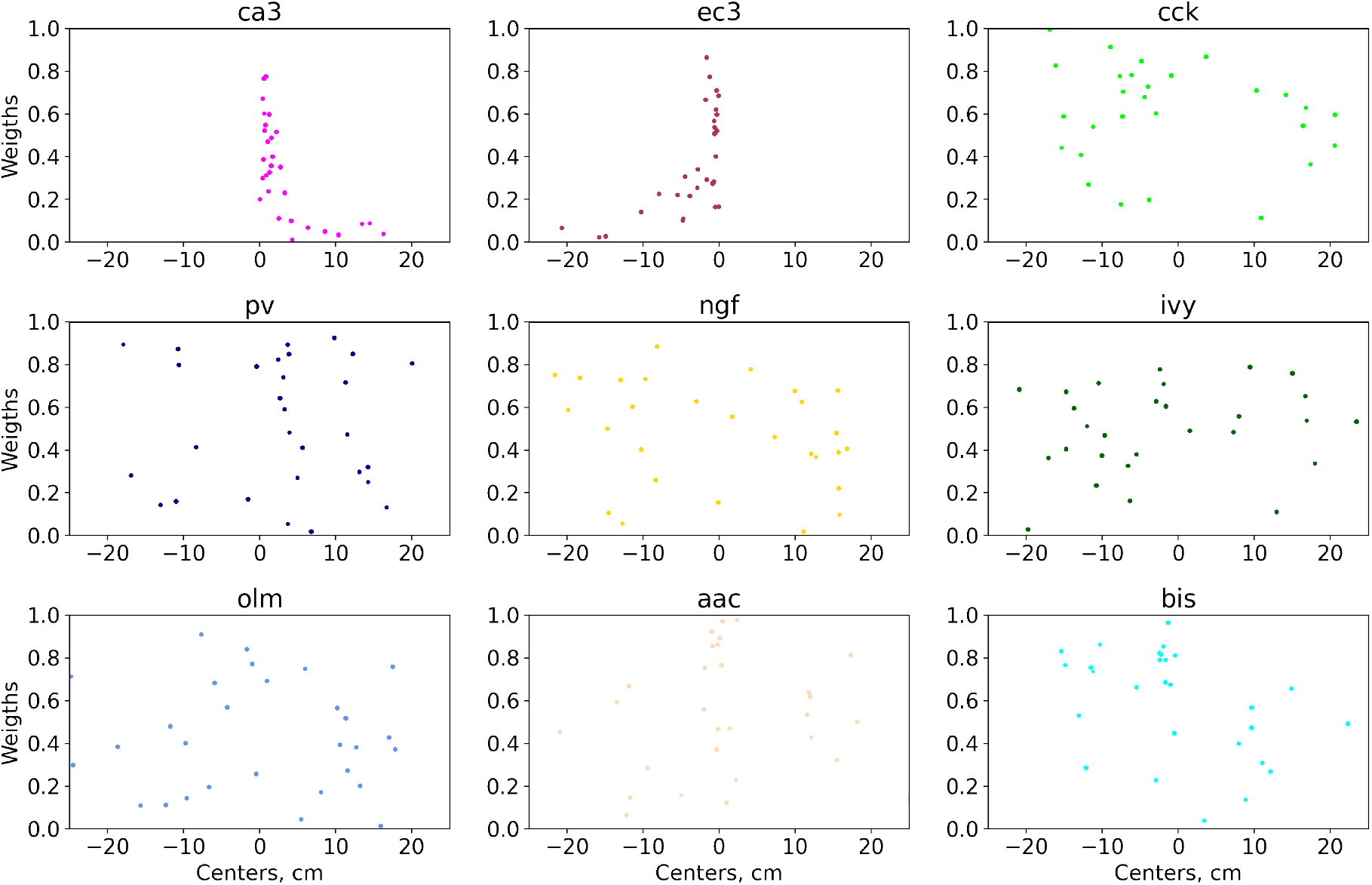
Plots are similar to Fig. 3. Optimization results for random sets of parameters. Non-optimized parameters are selected similar to Fig. 3. 30 sets of parameters have been studied. The spatial modulation of the inputs is limited, *σ* is no more than 8 cm.

## 4. Discussion

### Assumptions of the model

The differential evolution algorithm is well-established for finding global minima [25, 26], so the reliability of the conclusions depends entirely on the accuracy of the approximation of the inputs (eq. 3–5) and the output activity of a pyramidal neuron (eq. 7).

Our simulation results are based on several key assumptions. The most important assumption concerns the approximation of the spatial modulation of hippocampal neurons by a Gaussian function. The shape of the place fields of pyramidal neurons in field CA3 is well approximated by a Gaussian [36, 38]. The spatial activity of neurons in layer 3 of the EC can also be described using a Gaussian [39]. To describe the shape of the place fields for interneurons, we also chose Gaussian for two reasons. First, spatially modulated interneurons, regardless of class, have symmetrical place fields [22]. Second, interneurons within their place field are excited by unimodal input [20]. Both facts support the hypothesis that interneurons in the CA1 field inherit their spatial modulation from pyramidal cells. This hypothesis is supported by numerous studies in which, based on the correlation analysis of spike trains, the authors found many connections from pyramidal cells to interneurons [17, 40, 41, 42].

The second aspect of describing the inputs in our model was the approximation of theta modulation. We have used the von Mises function to generate the inputs. From the analysis of a large set of experimental data, it is known that more than 85% of CA3 neurons, EC layer 3, and CA1 field interneurons are modulated by the theta rhythm [10]. The degree of phase modulation R for all hippocampal neurons is in the range of 0.2 - 0.4; we used the value R = 0.35 as a median estimate. The anchoring phases for excitatory inputs and local interneurons (Table 1) are known from many experimental studies and are reliable [10, 12, 13, 43].

During the numerical solution of the optimization problem, multiple runs of the neuron model were required for different input configurations. Ferguson-Campbell neuron model was chosen by us as a compromise between biological plausibility and computational complexity [23].

### Comparison of our simulations with experimental results

Our results are in complete agreement with the experimental estimates presented by [20]. The authors recorded the activity of the pyramidal neurons of the CA1 field in freely-behaving animals and concluded a homogeneous level of inhibition inside and outside the place field. Grienberger and colleagues experimented with non-selective optogenetic depolarization of interneurons of all types, disinhibiting pyramidal cells. Grienberger et al’s conclusions are based on three experimental effects.

The disinhibition of pyramidal cells inside the field increases the spike rate more than outside the place field. If inhibition in the center of the place field were reduced, then the activation of pyramidal neurons would be insignificant. The somatic input impedance of the pyramidal cells was uniform inside and outside the place field in the control and experiments with the disinhibition. Typical autocorrelation somatic potential of the pyramidal neuron slightly increased (by about 10%) inside and outside the place field when the pyramidal cells were disinhibited [20]. Despite the absence of heterogeneity in inhibitory input, Grienberger found that non-selective suppression of interneurons reduced the strength of phase precession [20].

There are few studies on the contribution of individual groups of interneurons to phase precession. Our model predicts a large weight in the inhibitory input to axo-axonal neurons. An experimental study of axo-axonal cells shows that these neurons play an important role in controlling the activity of pyramidal neurons outside the place field. Optogenetic inhibition of axo-axonal neurons leads to the activation of pyramidal cells out of place and remapping in the CA1 field [44]. However, this study does not reveal the contribution of axo-axonal neurons to phase precession.

Royer and colleagues have shown that optogenetic suppression of PV neurons attenuates phase precession, shifting pyramidal activity to the descending theta phase when an animal is running into the place field [14]. Our model reproduces this effect, although in a weaker form (Fig. 4). It can be assumed that the effect obtained in [14] is also caused by partial suppression of axo-axonal neurons. Axo-axonal interneurons also express parvalbumin [43]; therefore, optogenetic inhibition also decreases the spike rate of these cells, which in turn provides the effect of attenuating phase precession. Another result presented in [14] is that the suppression of OLM neurons does not affect phase precession. This differs slightly from our estimates (Fig. 4).

We did not find experimental data in the literature on the role of other groups of interneurons in the formation of phase precession. The results presented in Fig. 4 make theoretical predictions. With optogenetic suppression of Ivy or CCK, the phase precession will weaken due to an increase in the activity of pyramidal neurons in the descending phase of the theta rhythm when an animal is running out of the place field. Suppression of neurogliaform neurons will lead to a shift in the activity of pyramidal neurons to the ascending phase of the theta rhythm when an animal is running into the place field. Despite the contradictory results, we expect that the activity of pyramidal neurons will not change significantly upon optogenetic suppression of bistratified cells. Notes, our model does not take into account interactions between interneurons, which may lead to slight differences between the experiment and theoretical predictions.

The hypothesis of homogeneous inhibition is supported by the data on the ratio of neural populations in the hippocampus. Interneurons make up only 9 to 20% of the neurons in the CA1 field [35]. At the same time, types of interneurons have very different numbers. For example, bistratified cells account for only 5.7% of all interneurons [35]. Therefore, it is most likely that interneurons do not encode information, but only modulate the activity of pyramidal cells by balancing inhibition at different phases of the theta rhythm. In this context, the distribution of interneuronal activity by theta rhythm phases is an important part of encoding information in the CA1 field.

### Comparison of our results with other models

A wide variety of theoretical models of phase precession has been proposed in the literature. The main ideas come down to interference of two inputs [37, 16, 45], asymmetric structure of one input [46, 47], short-term plasticity [48] or combinations of these ideas [49].

The role of interneurons in the formation of the place fields by pyramidal neurons has been little investigated in theoretical studies. The article [50] shows the importance of direct inhibition by PV basket neurons upon input from the CA3 field in stabilizing phase precession. A few other papers have generally noted the role of deceleration in the formation of place fields and phase precession [51, 52]. However, we did not find theoretical estimates of the structure of cell inhibition in the literature.

The hypothesis of interference of two oscillatory inputs for place cells in the CA1 field has received the most experimental support [17, 18]. Therefore, we took models based on this hypothesis [19, 37, 45] as a basis and developed this idea taking into account the latest experimental data on the parameters of modulation of interneuron firings by the theta rhythm.

### General Conclusion

Experimental studies of interneurons classes are to conduct. Only about 20% of neurons recorded in animals in free-behaving animals extracellularly are classified as interneurons [53]. This leads to the need to either do multiple experiments or use numerous electrodes to collect representative statistics, even without separating interneurons into classes.

The most promising methods for studying the interneurons are genetic modification methods: optogenetics and calcium fluorescence imaging. The reduction in the cost of these techniques leads to their availability for a greater number of laboratories. Attempts are being made to apply machine learning and big data techniques to identify firing patterns in electrophysiological extracellular cells corresponding to different types of interneurons [54]. To date, several databases have already been opened for analysis [53, 55, 56, 57].

We expect that in the coming years, interest in the role of interneurons in cognitive processes will increase due to the emergence of more opportunities for studying them. Theoretical studies of interneurons are important for determining the most promising directions for experimental research.

## Appendix

Equations for currents:

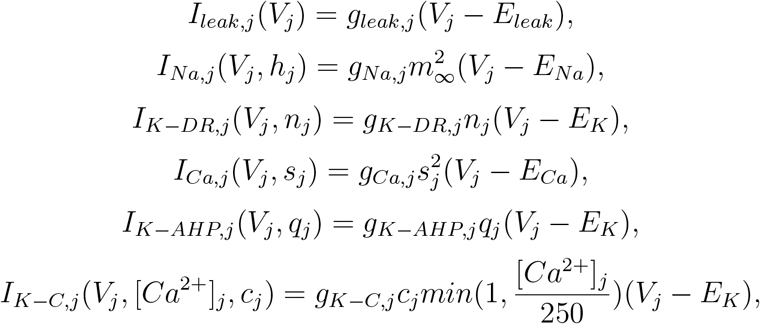

where *j* ∈ *S, D, S* and *D* denote the soma and dendrite of pyramidal neurons, respectively; E is the reversal potential of the currents (Table 3); *g* is the maximal conductance (Table 2). The equations and parameters for the synaptic currents (eq. 3) are given in the next section. All conductances are measured in the units *mS/cm*^2^.

**Table 2:** Maximal ionic conductances (*mS*/*cm*^2^)

| Compartment | $g_{Na,j}$ | $g_{Ca,j}$ | $g_{K-DR,j}$ | $g_{K-AHP,j}$ | $g_{K-C,j}$ | $g_{leak,j}$ |
| --- | --- | --- | --- | --- | --- | --- |
| Somatic (j = S) | 30.0 | 6.0 | 7.0 | 0.8 | 15.0 | 0.1 |
| Dendrite (j = D) | 0.0 | 5.0 | 0.0 | 0.8 | 5.0 | 0.1 |

**Table 3:** Values and units of reversal potentials, coupling parameters, and membrane capacitance

| Parameter | Unit | Parameter Value |
| --- | --- | --- |
| $E_{Na}$ | mV | 60 |
| $E_{Ca}$ | mV | 80 |
| $E_K$ | mV | -75 |
| $E_{leak}$ | mV | -60 |
| $g_C$ | $mS/cm^2$ | 1.5 |
| p |  | 0.5 |
| $C_m$ | $\mu F/cm^2$ | 3 |

The equations for intracellular calcium concentration [Ca2+] are:

Equations for calcium concentration

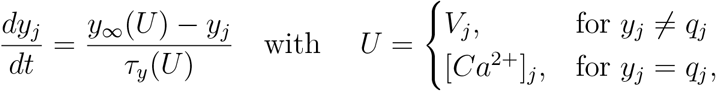

where *ϕ_Ca_* = 0.13 *mM* · *cm*^2^ · *nA*^−1^ is a scaling constant that converts the inward calcium current into internal calcium concentration, *β* = 0.075 *ms*^−1^.

The gating variables *h_j_*, *n_j_*, *s_j_*, *c_j_*, *q_j_*, *j* ∈ {*S, D*} are each governed by an equation of a form:

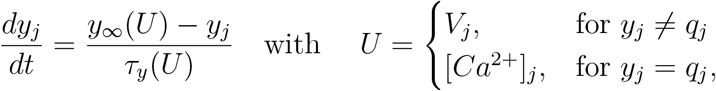

The associated steady state value and time constant are defined in the usual manner Equations for gate variables

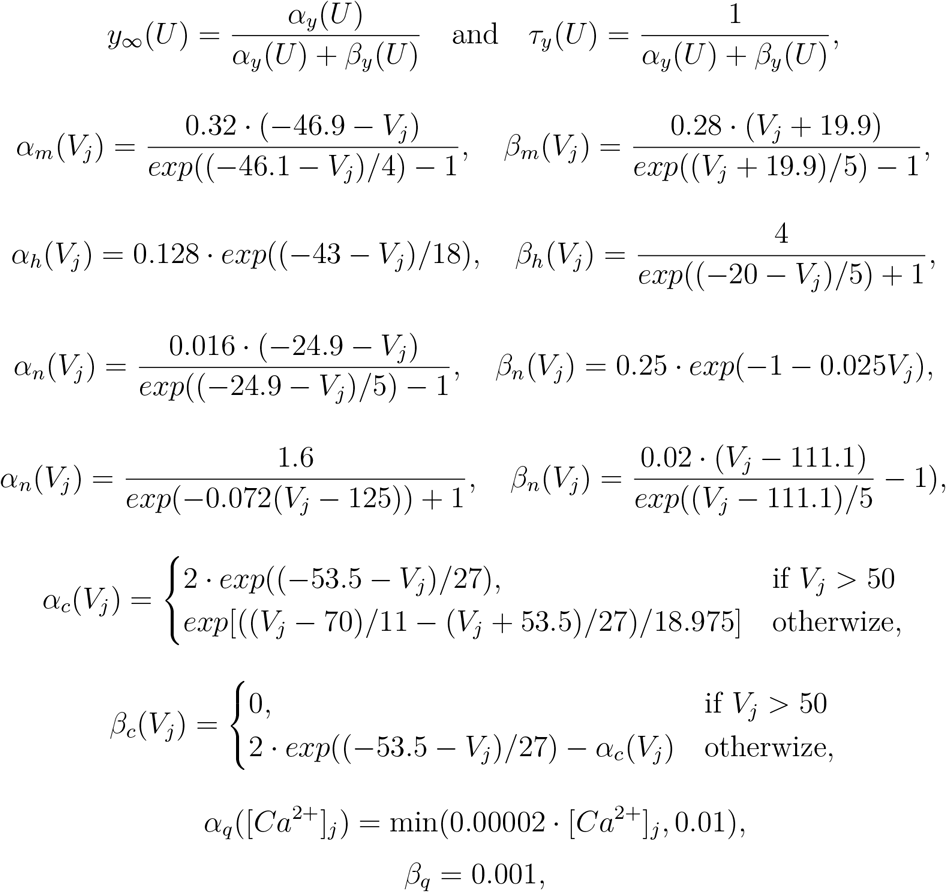

